## Supplementary figures and images for "Widespread conservation and lineage-specific diversification of genome-wide DNA methylation patterns across arthropods"

### Figure1_sup1

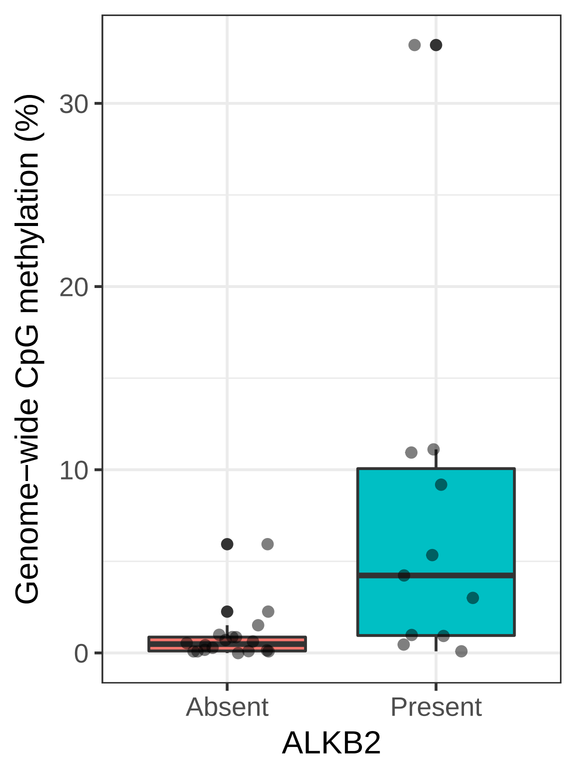

### Figure2_sup1

*Limulus polyphemus*

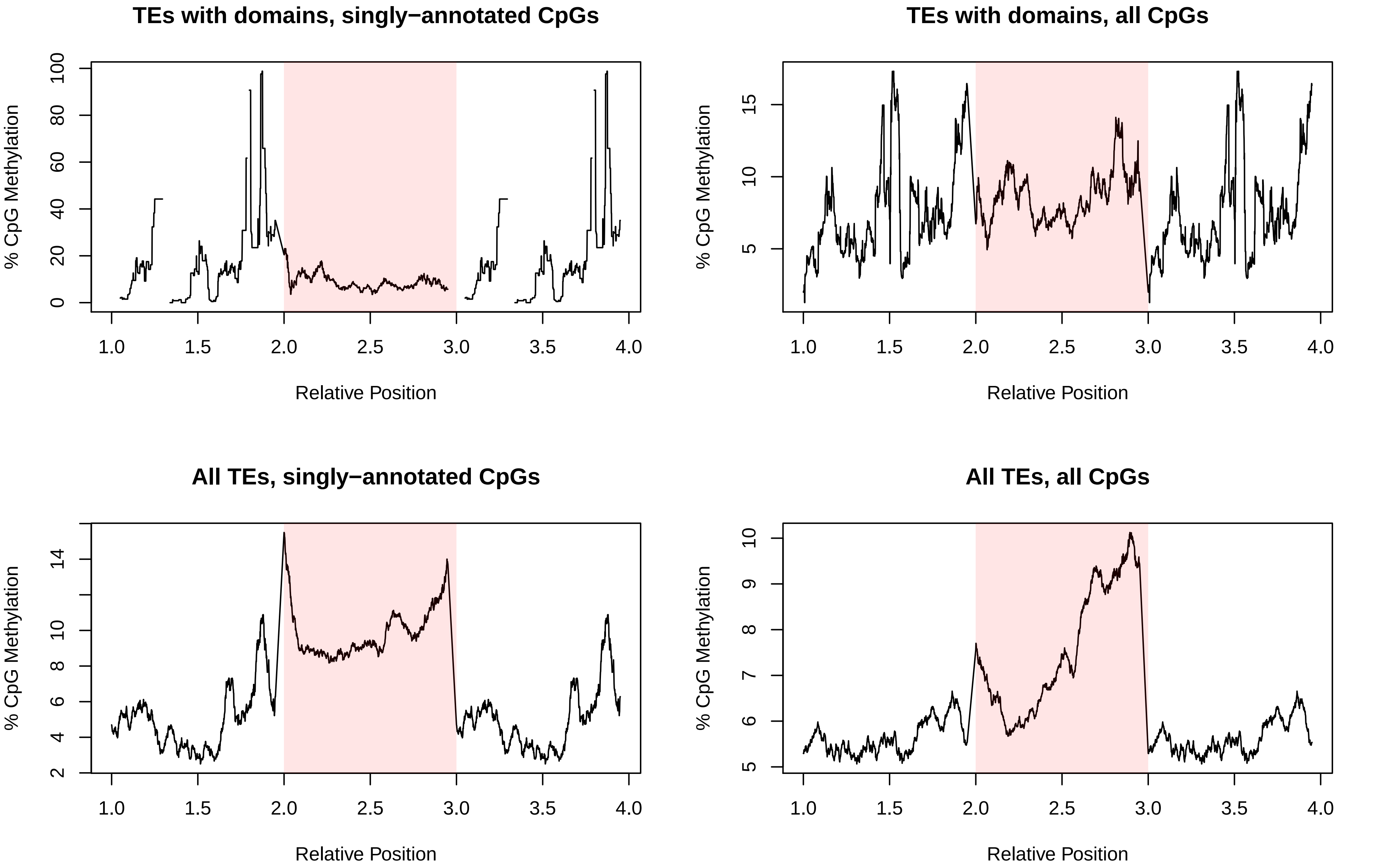

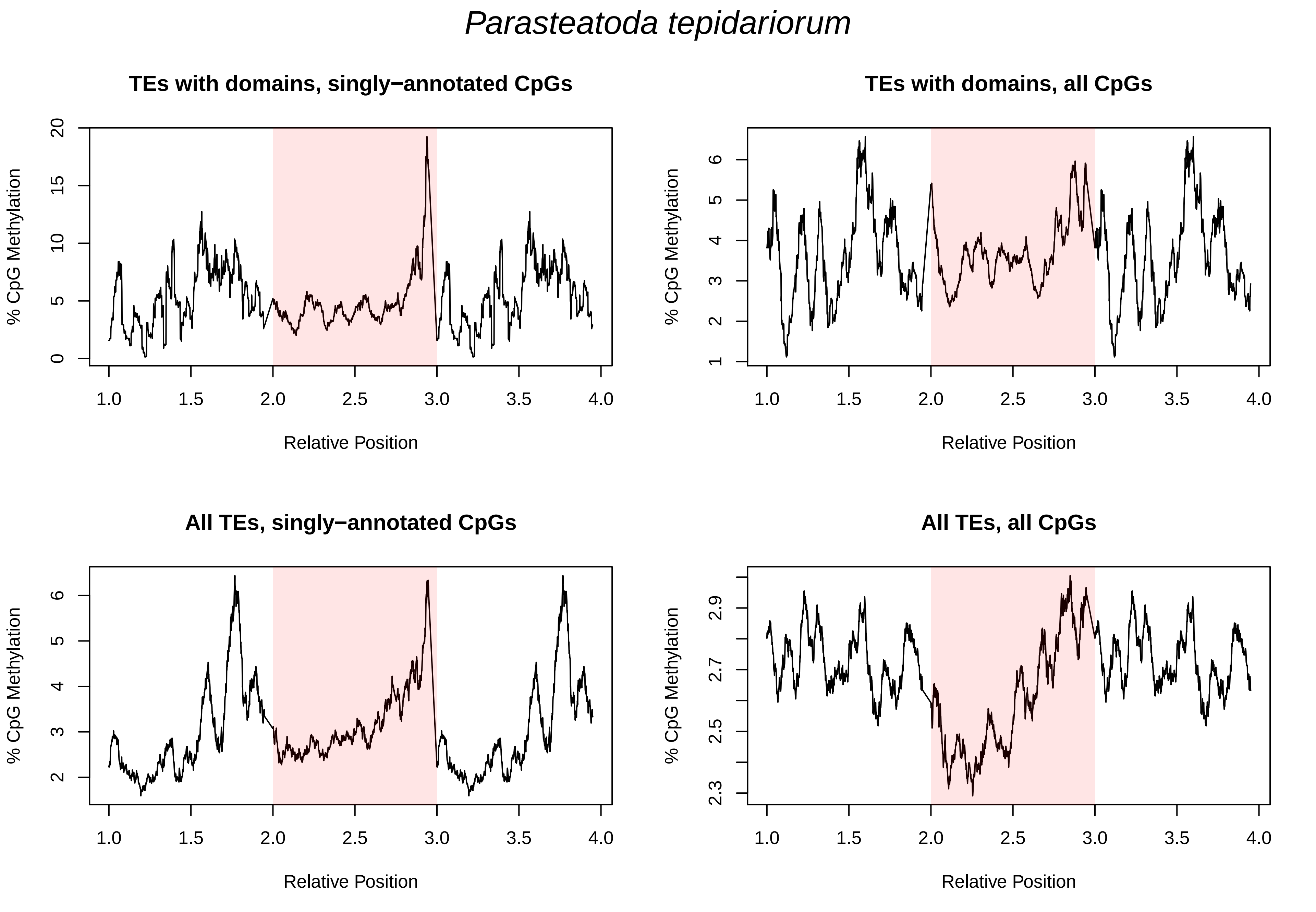

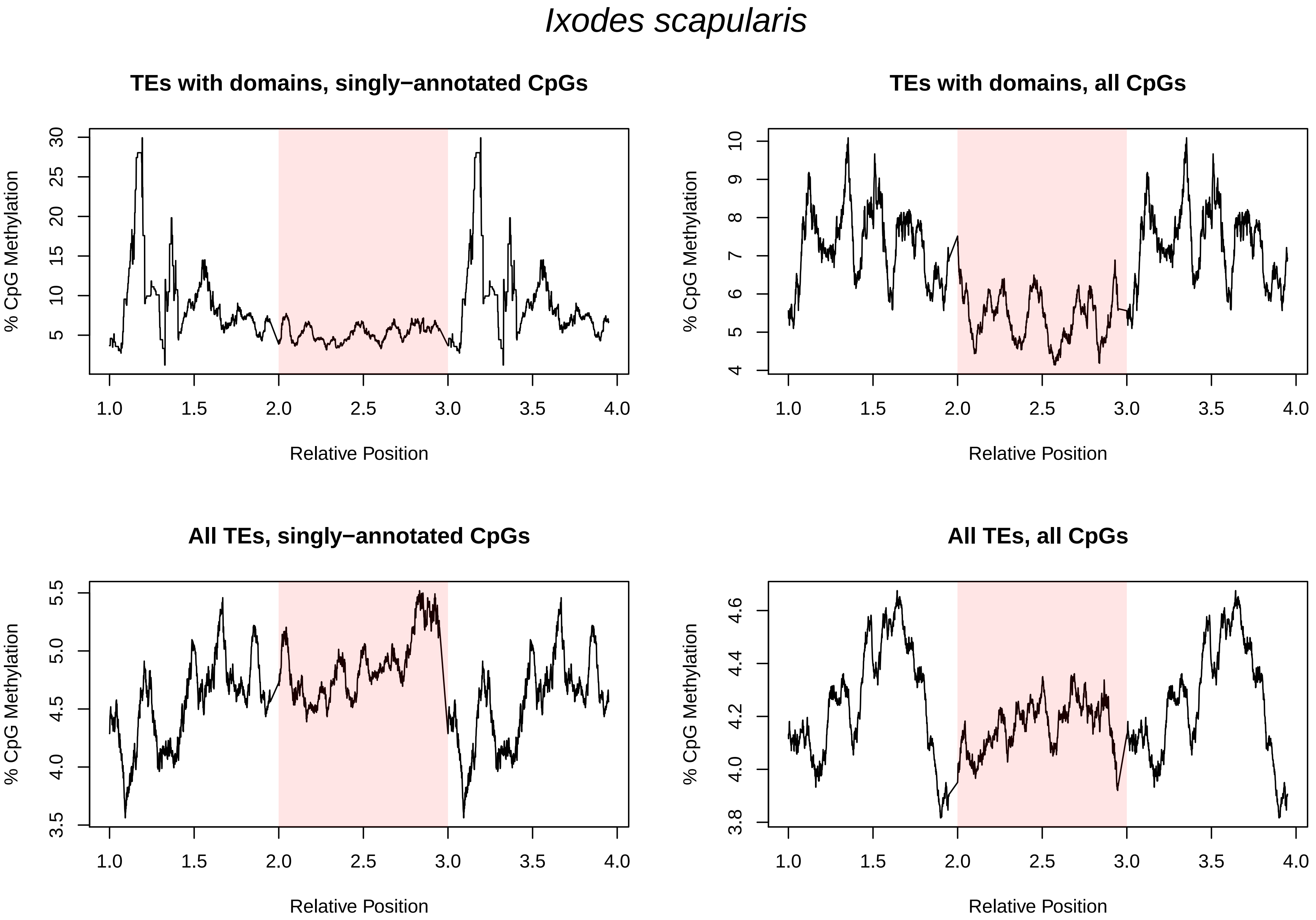

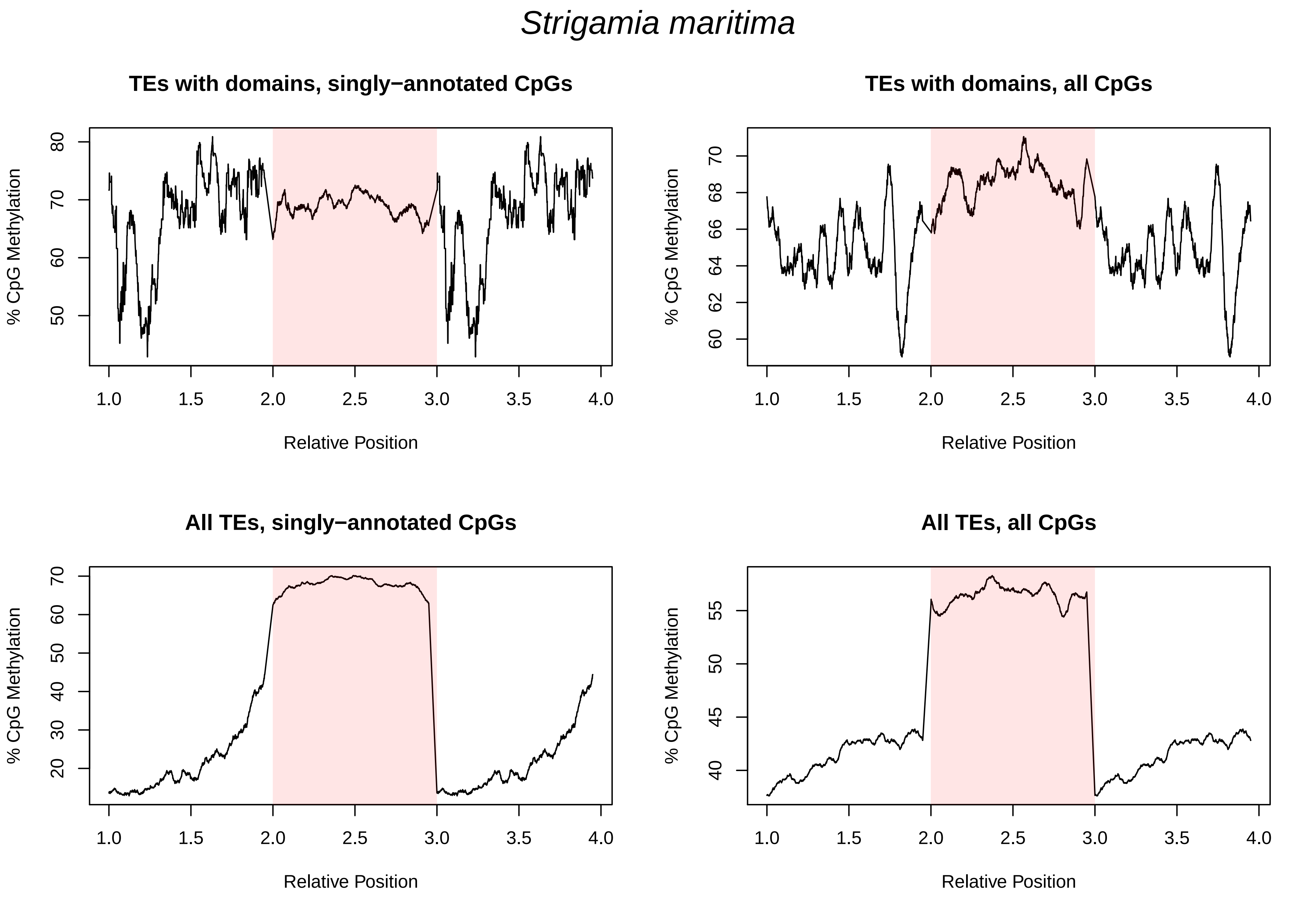

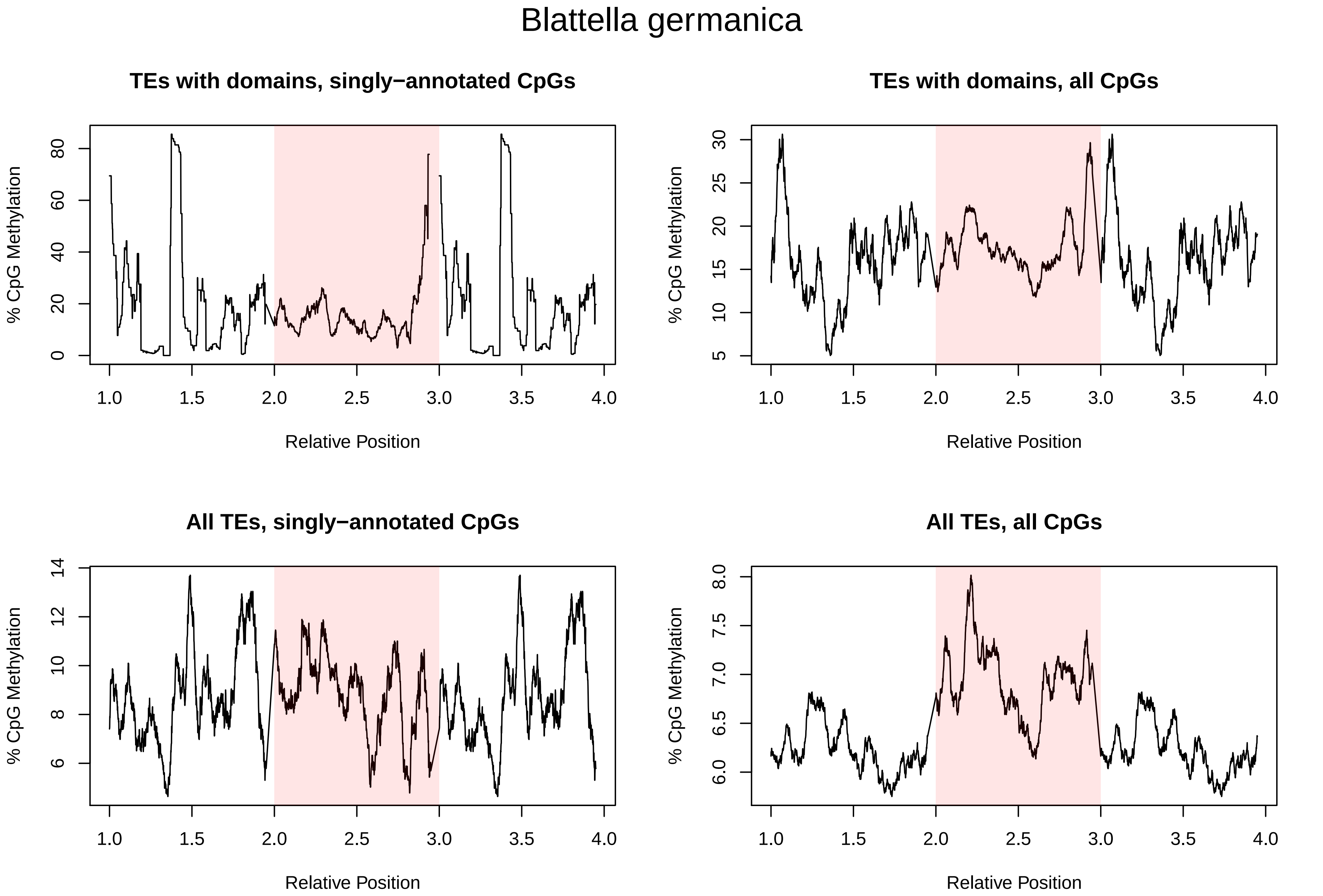

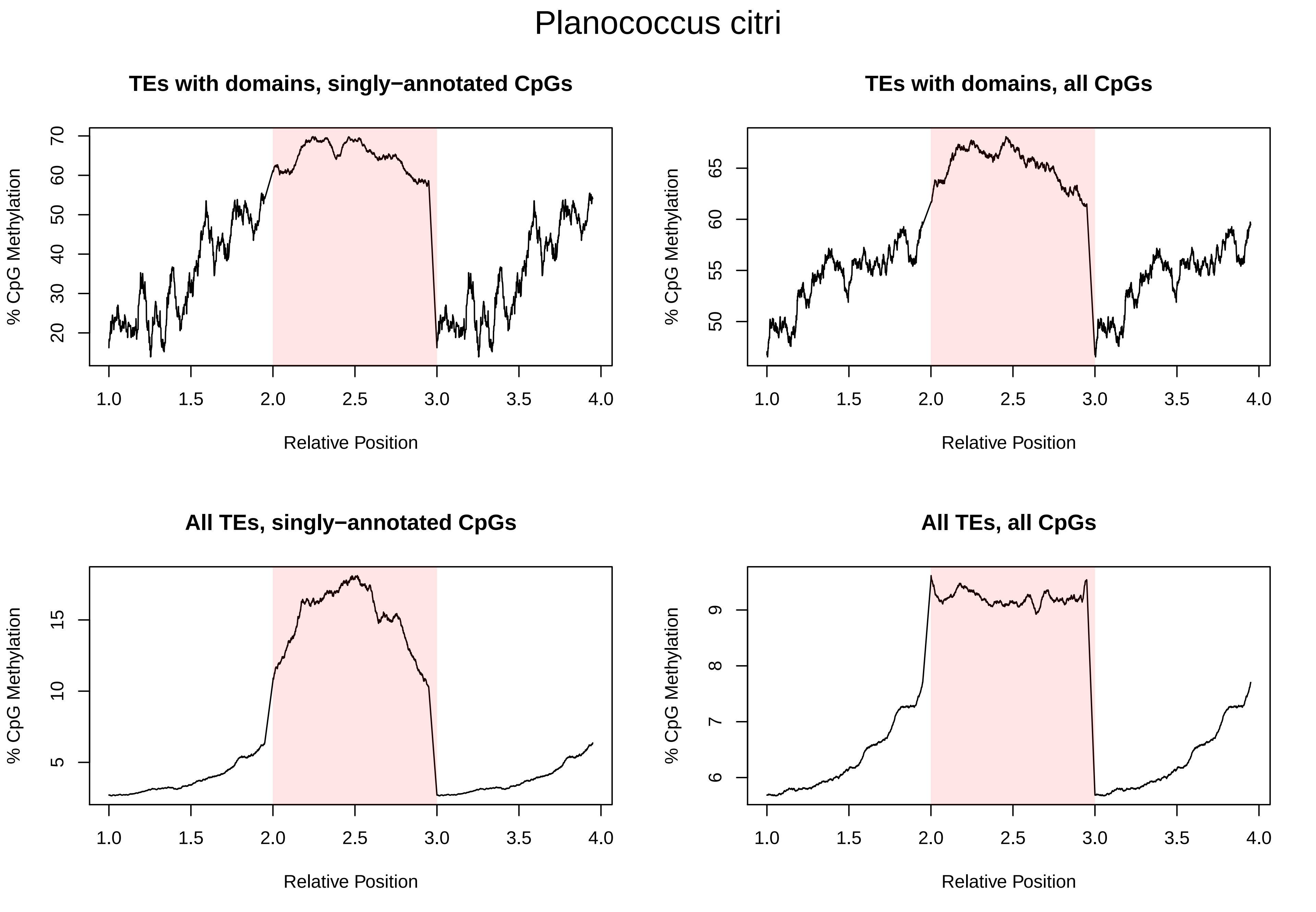

### Figure5_sup1

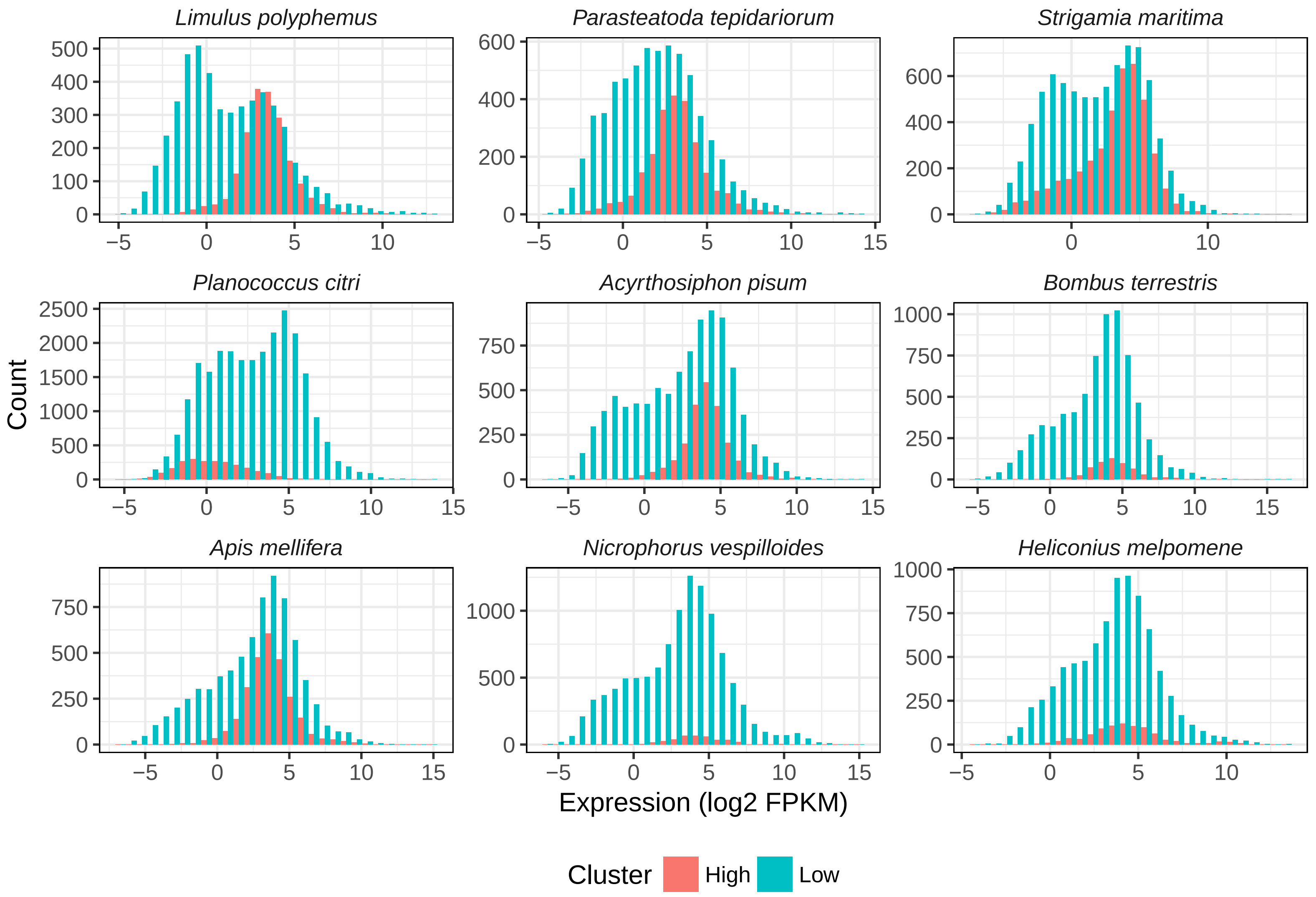

### Figure5_sup2

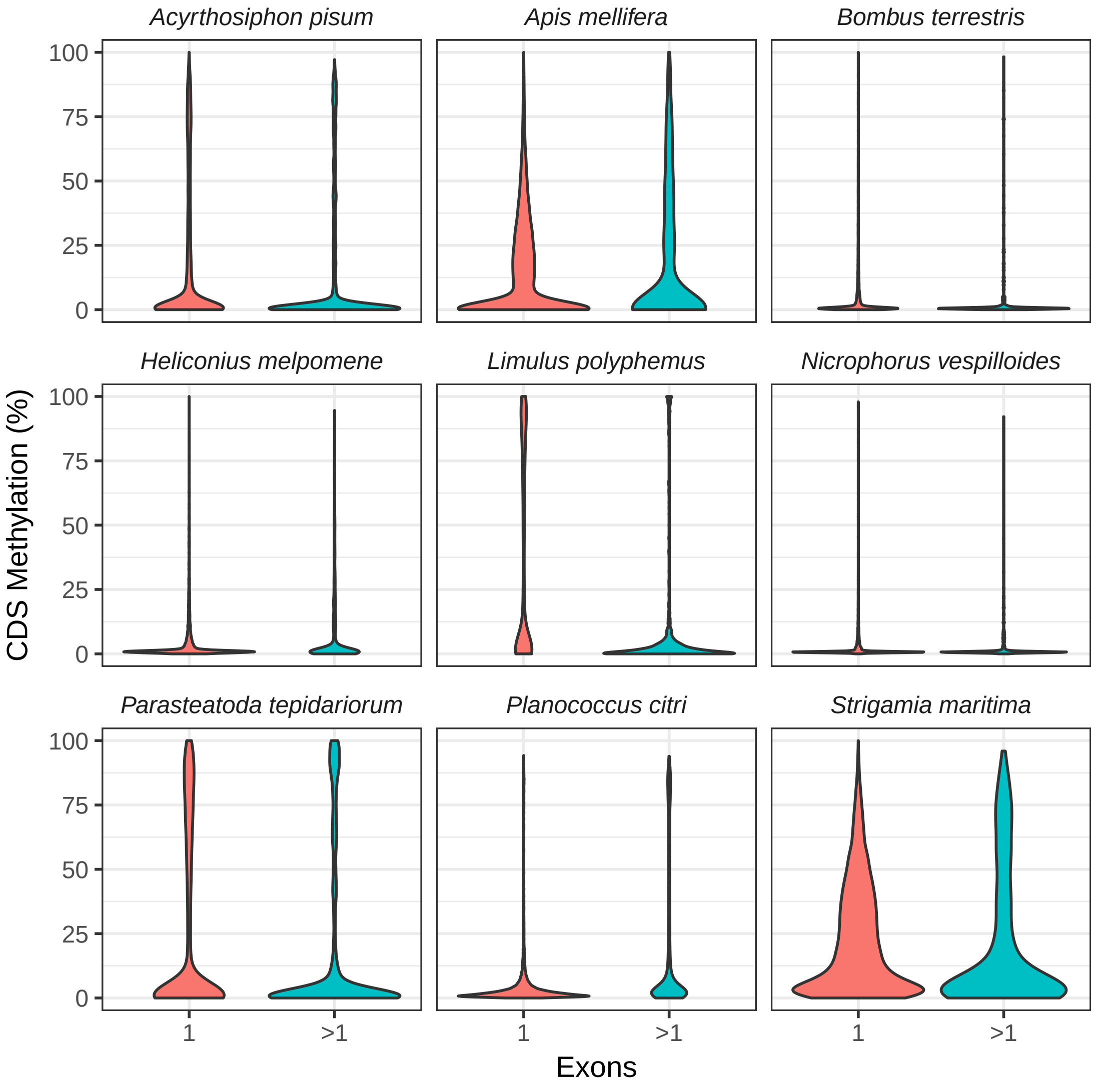

### Figure6_sup1

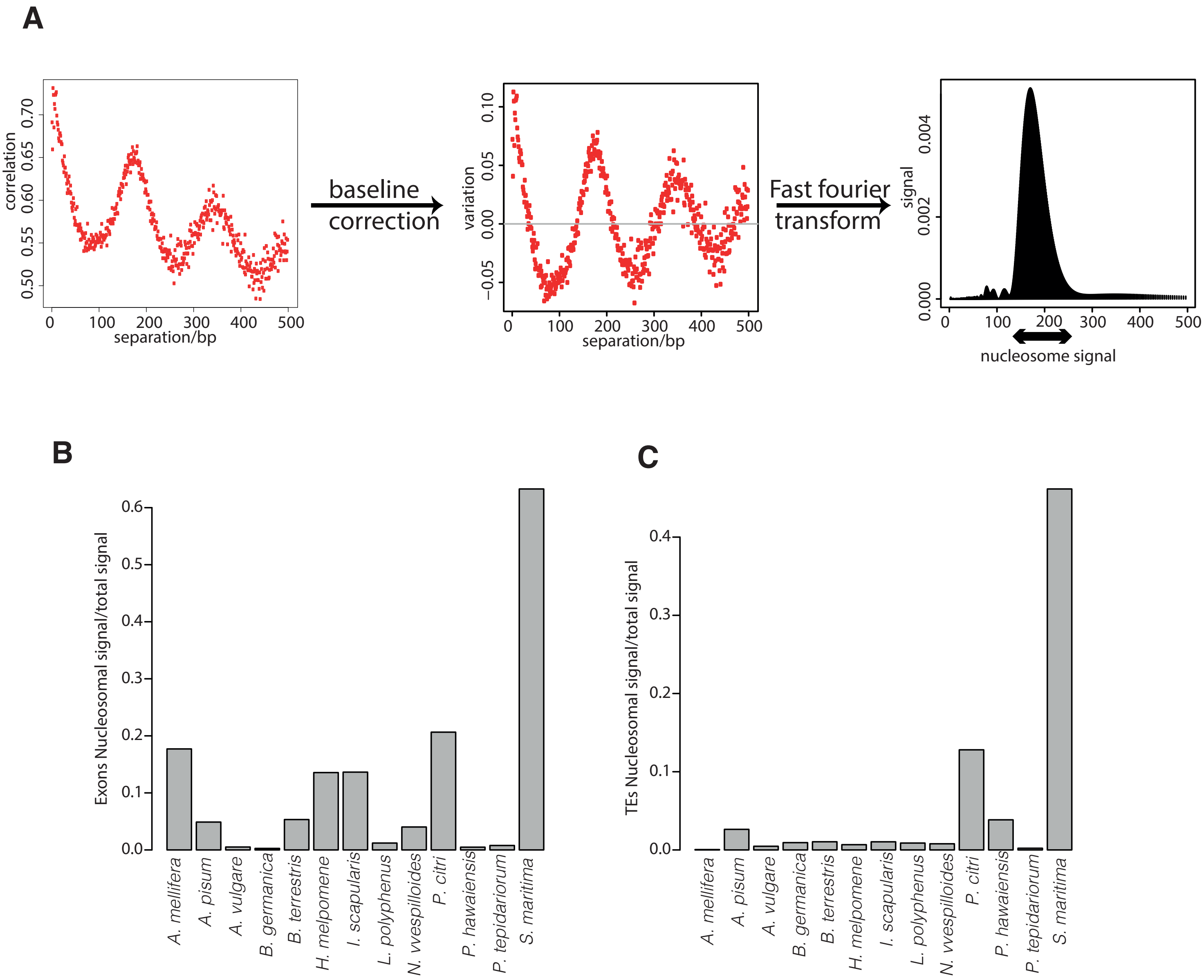

### Figure6_sup2

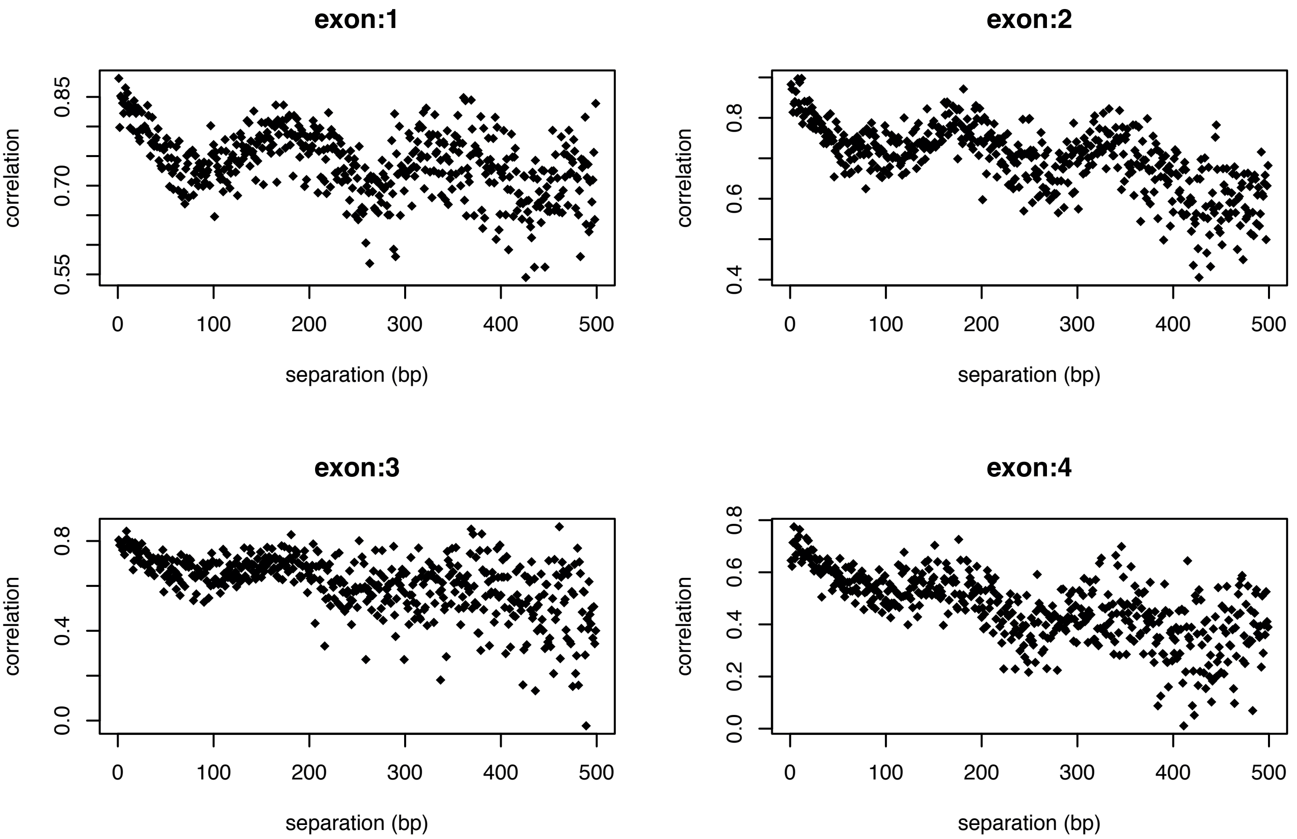

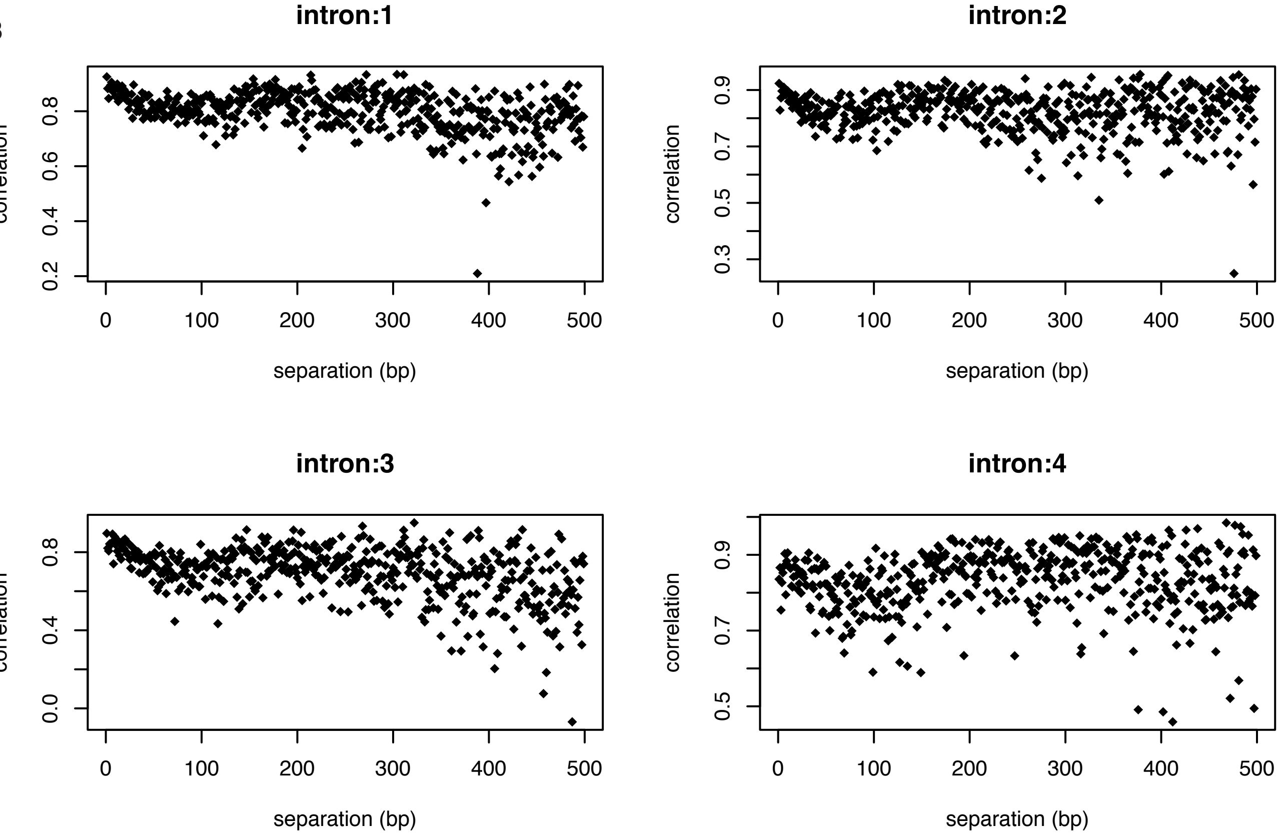
